# Identification of the Novel Pyroptosis-Related Gene Signature in Patients with Esophageal Adenocarcinoma

**DOI:** 10.1101/2021.07.05.451093

**Authors:** Ruijie Zeng, Shujie Huang, Zewei Zhuo, Huihuan Wu, Weihong Sha, Hao Chen

**Affiliations:** Department of Gastroenterology, Guangdong Provincial People’s Hospital, Guangdong Academy of Medical Sciences, Guangzhou 510080, China; Shantou University Medical College, Shantou 515041, China; Department of Thoracic Surgery, Guangdong Provincial People’s Hospital, Guangdong Academy of Medical Sciences, Guangzhou 510080, China

**Keywords:** Pyroptosis, Esophageal adenocarcinoma, Prognosis, Tumor microenvironment

## Abstract

Esophageal adenocarcinoma (EAC) is a highly malignant type of digestive tract cancers with a poor prognosis despite therapeutic advances. Pyroptosis is an inflammatory form of programmed cell death, whereas the role of pyroptosis in EAC remains largely unknown. Herein, we identified a pyroptosis-related five-gene signature that was significantly correlated with the survival of EAC patients in The Cancer Genome Atlas (TCGA) cohort and an independent validation dataset. In addition, a nomogram based on the five-gene signature was constructed with novel prognostic values. Moreover, the genes in the pyroptosis-related signature, *CASP1, GSDMB, IL1B, PYCARD*, and *ZBP1*, might be involved in immune response and regulation of the tumor microenvironment. Our findings indicate that the five-gene signature provides insights into the involvement of pyroptosis in EAC progression, and is promising in the risk assessment as well as prognosis for EAC patients in clinical practice.

## Introduction

Esophageal cancer is one of the most common malignancies worldwide, accounting for approximately 604,100 new cases and 544,076 deaths per year over the world ^1^. Esophageal adenocarcinoma (EAC) and esophageal squamous cell carcinoma (ESCC) composite the principle histologic subtypes of esophageal cancer, in which the incidence of EAC in western countries has increased dramatically in the last decades ^2^. Despite therapeutic advances in surgery, radiotherapy, chemotherapy, and targeted drugs, the 5-year survival of esophageal cancer remains less than 20% ^3^. In consequence, biomarkers and effective models are urgently needed to predict the prognosis of EAC and provide insights into targeted therapy.

Pyroptosis is a pro-inflammatory form of regulated cell death, relying on the enzymatic activity of inflammatory proteases that belong to the caspase family ^4^. Pyroptosis is featured with swift plasma-membrane rupture and subsequent release of pro-inflammatory intracellular contents, which is distinct from apoptosis ^5^. Studies evaluating the role of pyroptosis in neurological, infectious, autoimmune, cardiovascular and oncologic disorders have been emerging in recent years ^6^. Activation of the canonical inflammasome pathway is the basis of pyroptosis, in which pattern-recognition receptors (PRRs), for example, Toll-like receptors (TLRs), nucleotide-binding oligomerization domain-like receptor (NLRs) and absent in melanoma 2 like-receptor (ALRs) recognize pathogen-associated molecular patterns (PAMPs) or nonpathogen-related damage-associated molecular patterns (DAMPs) to activate inflammasomes and facilitate caspase-1 activation ^7^. Direct activation of caspase-4/5/11 under lipopolysaccharide (LPS) is involved in the noncanonical pyroptosis pathway, which is independent of the inflammasome complex ^8^. The gasdermin (GSDM) family proteins serve as the main mediators of pyroptosis, which are proteolytically activated by proteases and induce the formation of plasma membrane pores, leading to cell swelling and lysis ^9, 10^. Due to the pivotal role of GSDM family proteins, pyroptosis is defined by some researchers as gasdermin-mediated programmed cell death ^11^.

The role of pyroptosis has been explored in various types of cancer. GSDME is silenced in the majority of cancer cells, whereas expressed in various normal tissues ^12^. The knockdown of GSDMD inhibits the proliferation of non-small cell lung cancer (NSCLC) cells by intrinsic mitochondrial apoptotic pathways ^13^. The knock-out of inflammatory vesicle-related genes *in vivo* demonstrates the tendency to develop colon cancers ^14^. Metformin induces GSDMD-mediated pyroptosis and serves as an alternative treatment for therapy-refractory ESCC. Moreover, the PLK1 kinase inhibitor BI2536 sensitizes ESCC cells to cisplatin by inducing pyroptosis.

However, despite research is emerging in ESCC, the role of pyroptosis in esophageal cancer remains largely unknown, and none of the previous publications have comprehensively evaluated the pyroptosis-related genes in EAC. Therefore, we performed a comprehensive evaluation of pyroptosis-related genes in EAD, in order to develop a pyroptosis-gene-based modality to predict the prognosis of the patients, and provide insights into the correlations between pyroptosis and tumor immune microenvironment.

## Methods

### Datasets

The RNA-sequencing (RNA-seq) data of 87 patients (78 with EAD; 9 normal samples) and the corresponding clinical information from The Cancer Genome Atlas (TCGA) database were retrieved on May 20^th^, 2021 (https://portal.gdc.cancer.gov/repository). The RNA-seq data and clinical features of the validation cohort were downloaded from the Gene Expression Omnibus (GEO) database (https://www.ncbi.nlm.nih.gov/geo/, ID: GSE13898). Compared to the followup time of the TCGA cohort, that of the GSE13898 cohort was shorter (up to 4.9 years).

### Identification of differentially expressed genes (DEGs) in pyroptosis-related gene set

The 58 pyroptosis-related genes were derived from prior literature and the Gene Ontology (GO) term pyroptosis (ID: GO0070269; Supplementary Table S1) ^7, 15, 16, 17^. Normalization of expression data to fragment per kilobase million (FPKM) values was performed in both datasets. The package “limma” was used to explore DEGs with the threshold of *P* value < 0.05 ^18^. Protein-protein interaction (PPI) networks were created by Search Tool for the Retrieval of Interacting Genes (STRING) and the “igraph” package ^19, 20^.

### Development and validation of the pyroptosis-correlated gene prediction model for prognosis

Cox regression analysis was employed to evaluate the value of pyroptosis-related genes for prognosis. The DEGs were identified for further analysis. The LASSO Cox regression analysis was employed to construct a refined model for prognosis using the R package “glmnet” ^21^. The calculation of the risk score was performed using the following formula: 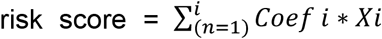 (Coef i indicates the coefficient, and Xi indicates the gene expression levels after standardization). The EAC patients were classified into low- and high-risk groups based on the median risk score, and Kaplan-Meier analysis was used to compare the overall survival (OS) between the two groups. Principal component analysis (PCA) was used to assess the separability of the two groups by the “prcomp” function. The R packages “survival”, “survminer”, “timeROC” and “riskRegression” were utilized for receiver operating characteristic (ROC) curve graphing and area under curve (AUC) calculation for 1, 2 and 5 years ^22, 23, 24, 25^. A nomogram model with clinical features including stage and risk score was constructed by the R packages “rms”, “foreign” and “survival” ^22, 26, 27^. The calibration curve and detrended correspondence analysis (DCA) were performed using the “rms” package ^26^. An EAC cohort (GSE13898) from the GEO database was used for validation, and the risk score was calculated by the same methods described above to divide the cohort into two subgroups (low risk and high risk).

### Prognostic analysis of the variables

Clinical data (gender and stage) was extracted of patients in the TCGA cohort and the GSE13898 cohorts. Variables including gender, stage and risk score were analyzed in the regression model by univariate and multivariate Cox regression analysis.

### Enrichment analysis

Patients with EAC in the TCGA cohort were divided into two groups based on the median risk score. The DEGs between the low- and high-risk groups were extracted by |log2FC| ≥ 1 and *P* value < 0.05. GO and Kyoto Encyclopedia of Genes and Genomes (KEGG) pathway enrichment was performed by the R package “clusterProfiler”, and the results were visualized using the “GOplot” package ^28, 29^.

### Tumor microenvironment analysis

The Tumor Immune Estimation Resource (TIMER) database (https://cistrome.shinyapps.io/timer/) was utilized to assess the correlation between tumorinfiltrating immune cells and expressions of selected genes ^30^. Estimation Resource (TIMER) was used to compare the immune scores of the four subtypes. The CIBERSORT algorithm was used to further explore the composition and differences in the fraction of 22 immune cell types between two subgroups classified by risk scores ^31^.

### Statistical analysis

Statistical analyses were performed by R (version 4.1.0). Student’s t-test was applied to compare the differences in gene expression between tumor and normal tissues, while categorical variables were compared using Pearson chi-square test. The OS of patients between low- and high-risk groups were compared by the Kaplan-Meier method with log-rank test. The Cox regression analysis was performed to evaluate the independent prognostic factors for survival. The Wilcoxon test was used to compare the immune cell infiltration between groups.

## Results

### Identification of DEGs between EAC and normal tissues

The expression levels of 58 pyroptosis-correlated genes were examined in the TCGA data of 78 EAC and 9 normal tissues. Ten DEGs were identified (|log2FC| ≥ 1 and *P* value < 0.05), and all of them (*CASP1, CASP5, GSDMB, GZMB, IL1B, NLRP6, PYCARD, TNF, TREM2, ZBP1*) were upregulated in the tumor group. The expression profiles of DEGs were demonstrated in **Fig. 1A** (red color represents higher expression level; blue color represents lower expression level). **Fig. 1B** showed the correlation network of DEGs in the TCGA data. The PPIs of DEGs were presented in **Fig. 1C**, in which the interaction score was set as 0.4.

**Figure 1.**
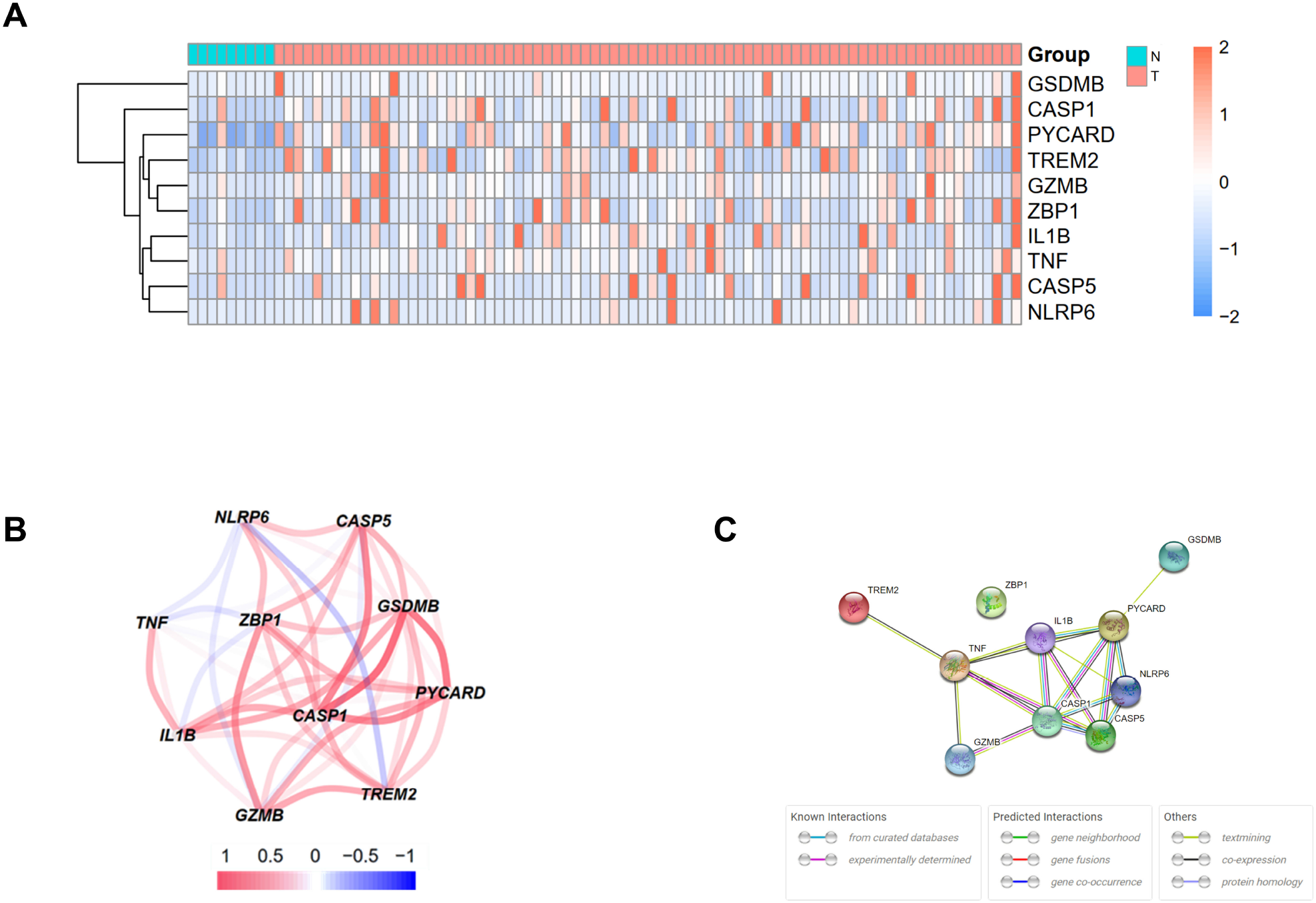
Expressions of 10 differentially expressed pyroptosis-related genes and interaction. **A.** Heatmap of gene expression between the tumor (T, red color) and normal (N, blue color) group. Higher expression: red color. Lower expression: blue color. **B.** Correlation network of 10 differentially expressed pyroptosis-related genes (red: positive correlation; blue: negative correlation; deeper color indicates higher strength of relevance). **C.** Protein-protein interaction (PPI) network of proteins encoded by selected genes.

### Construction of prognostic model based on DEGs

A total of 65 EAC patients with available survival data were included in our study. Univariate Cox regression analysis was initially performed to assess the prognostic value of DEGs **(Fig. 2A)**. Among them, 6 genes (*CASP1, CASP5, GSDMB, IL1B, PYCARD and ZBP1*) were with the *P* value < 0.2, and higher expressions of *CASP1, CASP5, IL1B* were associated with increased risk (HR > 1), while upregulated expressions of *GSDMB, PYCARD, ZBP1* were correlated with lower risk (HR < 1). Subsequently, LASSO Cox regression analysis retrieved 5 genes for prognostic model construction based on the optimum λ value (**Fig. 2B, 2C**). The calculation of risk score was as follows: Risk score = (0.042 * exp*CASP1*) + (−0.025 * exp*GSDMB*) + (0.021 * expIL1B) + (−0.037 * exp*PYCARD*) + (−0.243 + exp*ZBP1*). According to the calculated median risk score, 65 patients were divided into two groups (32 in the high-risk group and 33 in the low-risk group), and the clinical information was shown in **Fig. 3A**. The PCA analysis illustrated that patients were well divided into two clusters (**Fig. 3B**). The distributions of risk score and survival time were shown in **Fig. 3C, 3D**. The OS of the high-risk group was significantly worse than that of the low-risk group (*P* = 0.0012, **Fig. 3E**). ROC analysis of the risk model indicated that the AUC for 1-year, 2-year and 5-year survival was 0.708, 0.815 and 0.952, respectively (**Fig. 3F, 3G, 3H**). Both of the univariate and multivariate Cox regression analyses showed that the pyroptosis-related gene signature independently predicted the prognosis of EAC patients (**Fig. 3I, 3J**).

**Figure 2.**
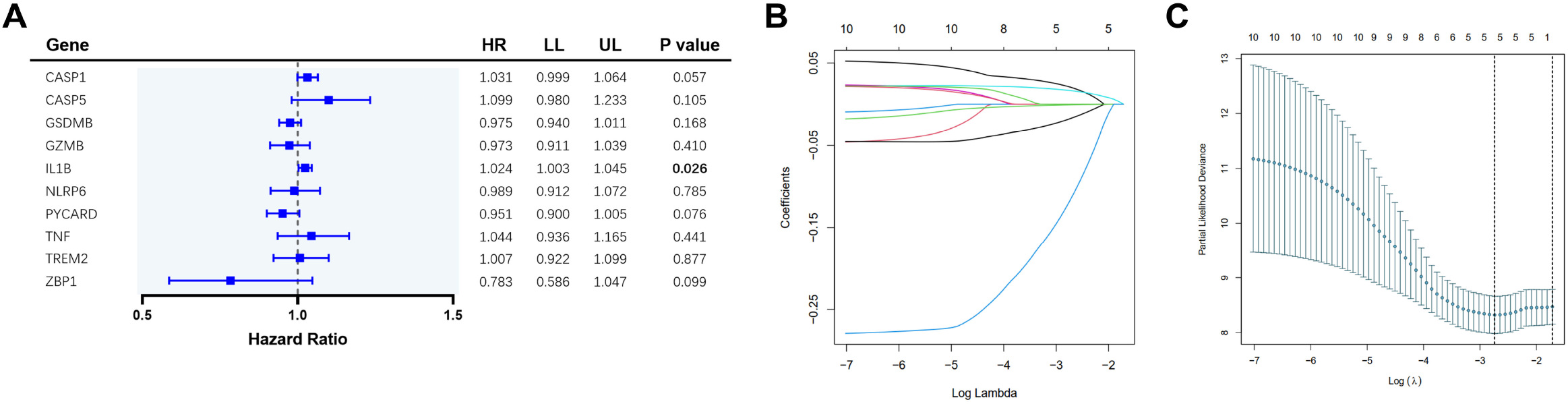
Construction of pyroptosis-related gene risk signature in TCGA cohort. **A.** Univariate cox regression analysis of overall survival (OS) for the selected genes. **B.** LASSO regression of the 10 selected genes. **C.** Cross-validation for tuning the parameter selection in lasso regression.

**Figure 3.**
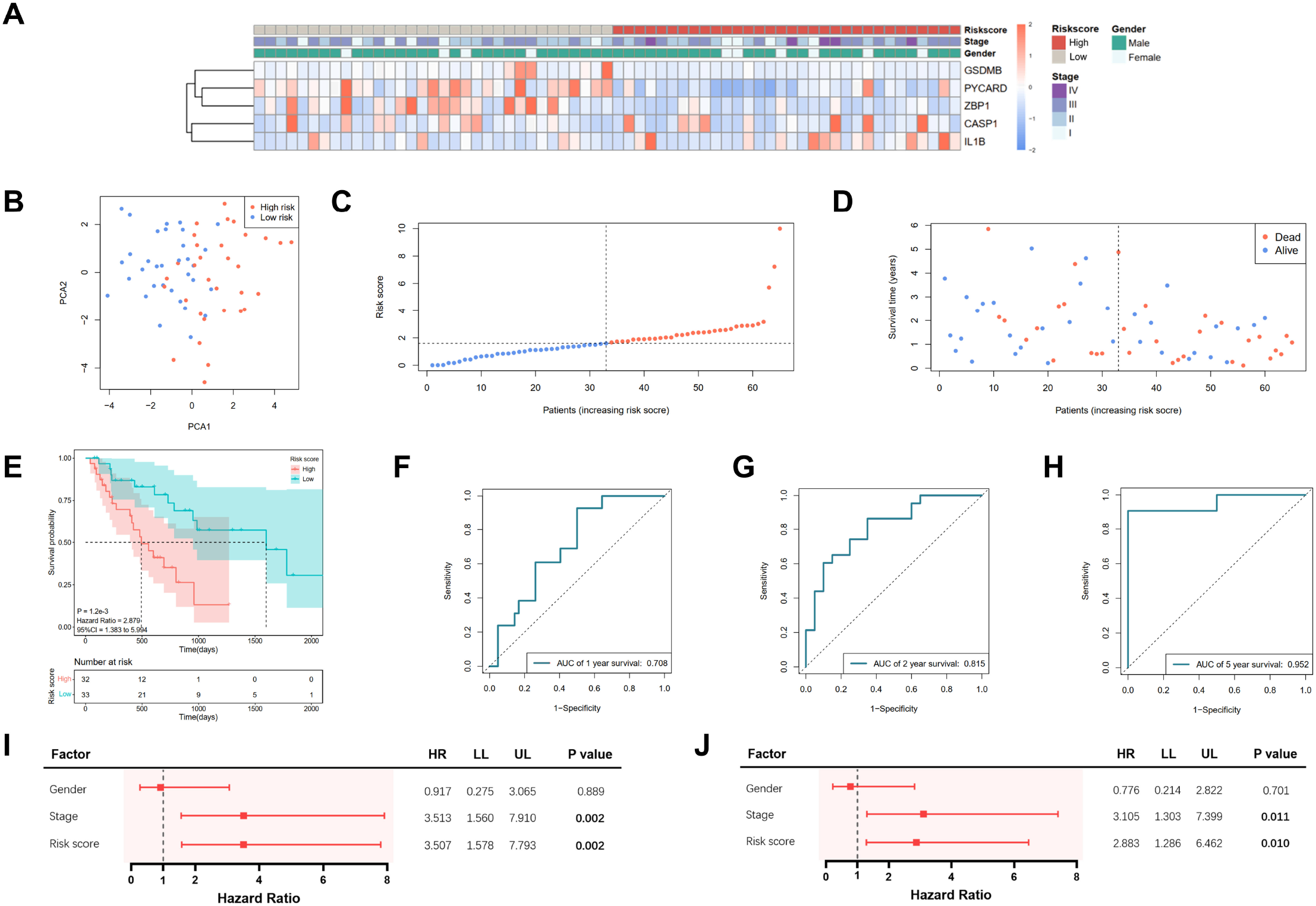
Prediction of prognosis using the pyroptosis-related five-gene signature in TCGA cohort. **A.** Heatmap of 5 selected gene expressions with clinical features ordered by risk score (red: higher expression; blue: lower expression). **B.** Principal component analysis (PCA) of the risk groups. **C.** Distribution of patients according to risk scores. **D.** Survival time and status of patients. **E.** Kaplan-Meier curves for the survival of patients in the low- and high-risk groups. **F, G, H.** Receiver operating characteristic (ROC) curve for 1-, 2- and 5-year survival of patients. **I.** Univariate Cox analysis. **J.** Multivariate Cox analysis.

### Verification of the gene signature by the external dataset

Information of 60 EAC patients from the GSE13898 dataset of GEO with available survival data was used for validation of the 5-gene signature. The patients were subdivided into the low- and high-risk group respectively as described above. PCA illustrated well separation of patients between the two groups (**Fig. 4A**). The distribution of risk score and survival time was demonstrated in **Fig. 4B, 4C**. Patients in the low-risk group were with significantly higher survival rates than those in the high-risk group (*P* = 0.003; **Fig. 4D**). According to the ROC curve, the 1-year and 2-year survival prediction models were with the AUC of 0.678 and 0.663 (**Fig. 4E, 4F**), respectively, while the 5-year survival prediction model could not be generated due to insufficient data. The risk score in our model could also serve as an independent prognostic factor in the validation cohort (**Fig. 4G, 4H**).

**Figure 4.**
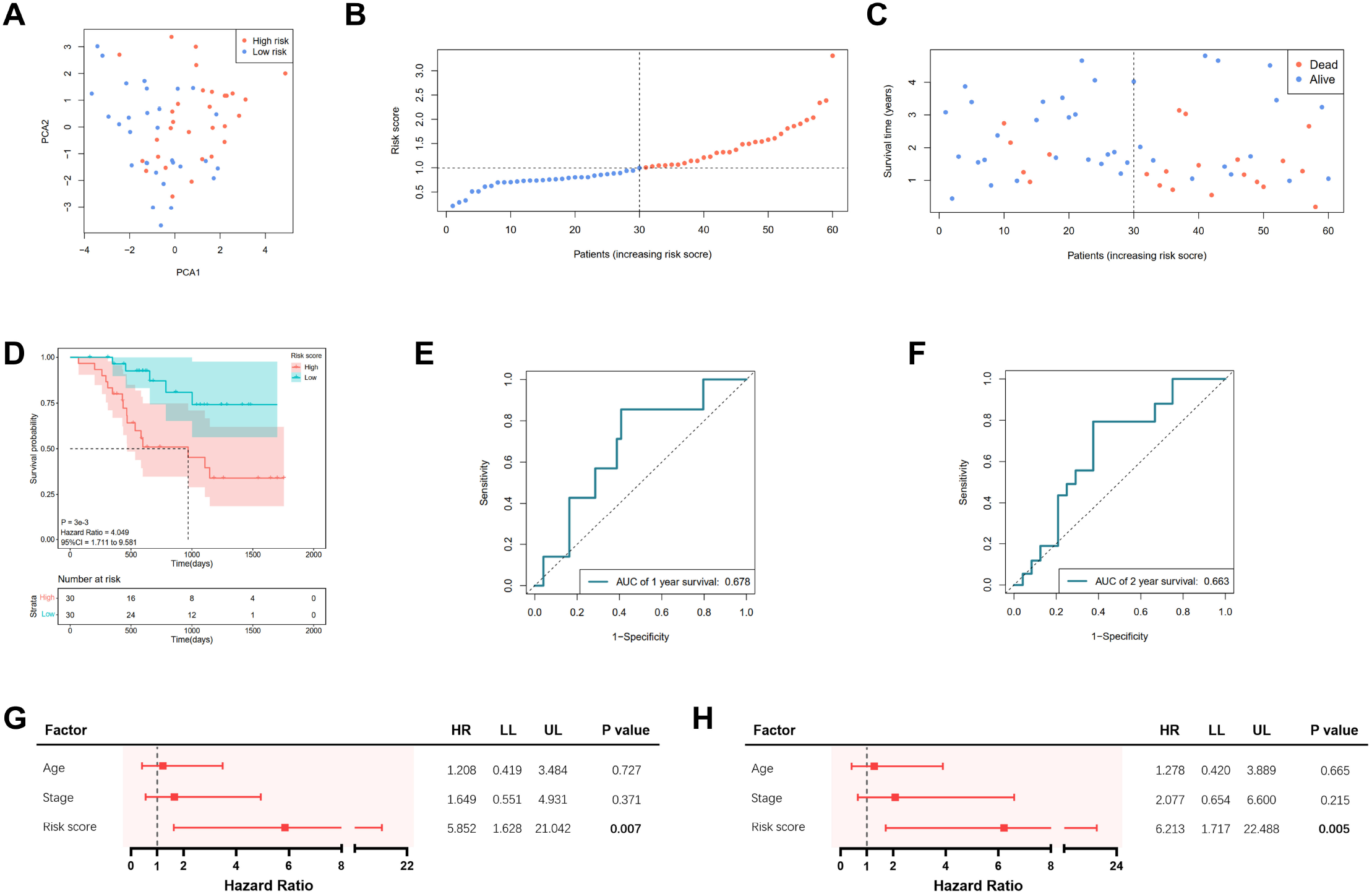
Validation of the pyroptosis-related five-gene signature in GSE13898 cohort. **A.** Principal component analysis (PCA) of the risk groups. **B.** Distribution of patients according to risk scores. **C.** Survival time and status of patients. **D.** Kaplan-Meier curves for the survival of patients in the low- and high-risk groups. **E, F.** Receiver operating characteristic (ROC) curve for 1-, 2- and 5-year survival of patients. **G.** Univariate Cox analysis. **H.** Multivariate Cox analysis.

### Construction of nomogram based on the gene signature and clinical data

In order to more precisely predict the prognosis of EAC patients, TNM stage was used to construct a nomogram model as shown in **Fig. 5A**. The AUCs of the nomogram for predicting 1-year, 2-year and 5-year survival were 0.722, 0.884, 1.00, respectively (**Fig. 5B, 5C, 5D**). The calibration curve indicated an ideal prediction of the nomogram (**Fig. 5E**). **Fig. 5F** showed that when the nomogram-predicted probability was ranged from 15% to 80%, the nomogram provided extra value relative to the treat-all-patients scheme or the treat-none scheme.

**Figure 5.**
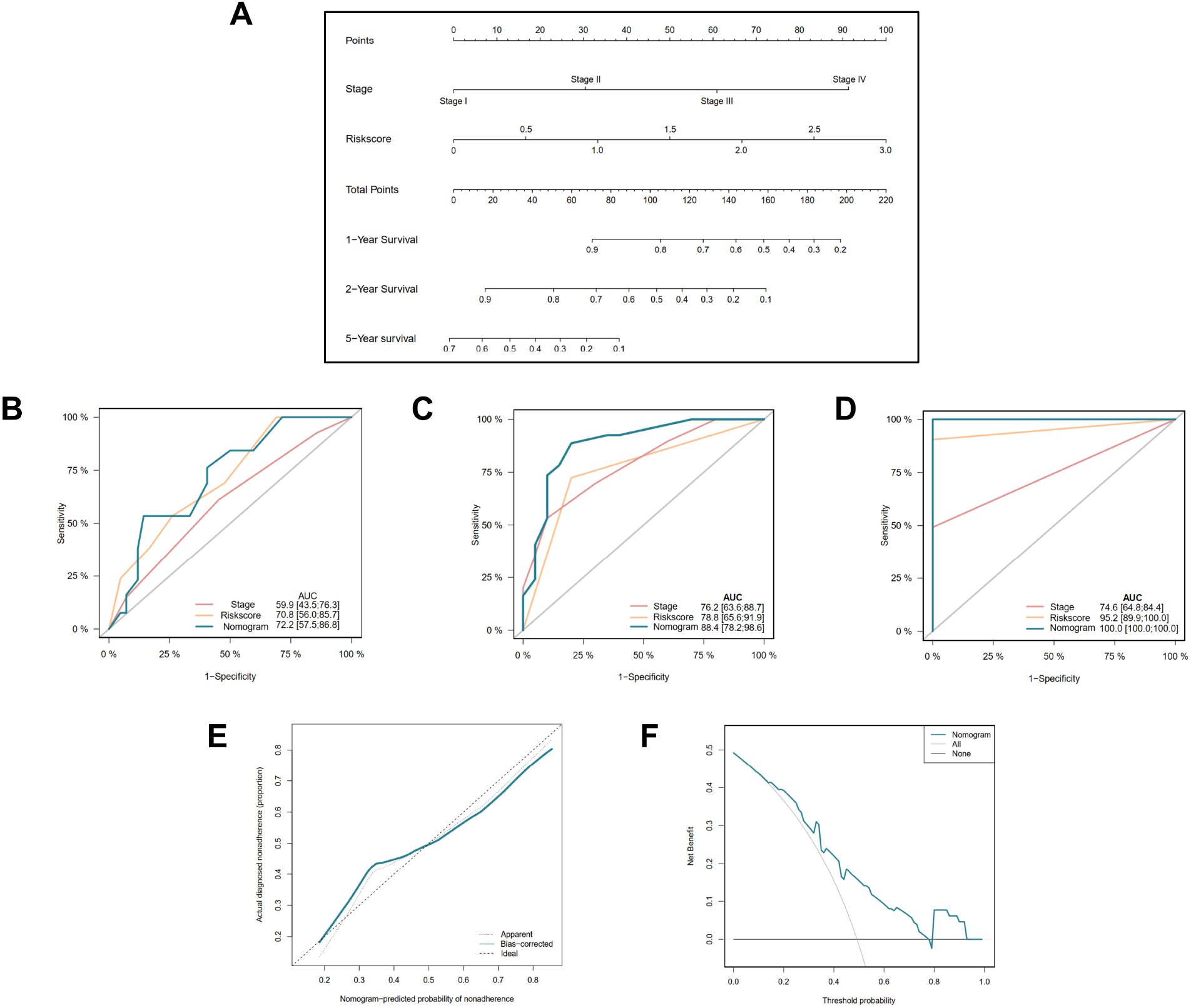
Construction of nomogram based on the pyroptosis-related five-gene signature. **A.** Nomogram for predicting 1, 2, 5-year survival of EAC patients. **B, C, D.** Receiver operating characteristic (ROC) curve evaluating the efficiency of nomogram for 1-, 2- and 5-year survival of patients. **E.** Calibration curve of nomogram. **F.** Decision curve analysis (DCA) curve of the nomogram.

### Differential expression analysis

A total of 527 DEGs between the low- and high-risk groups were extracted according to the threshold described above. A total of 310 genes were downregulated in the high-risk group, while 217 genes were upregulated in the low-risk group (**Table S2**). On the basis of the DEGs, GO enrichment and KEGG pathway analyses were performed. The results from GO enrichment analysis demonstrated that the DEGs were mainly associated with the regulation of cytokine production, cytokine activity and humoral immune response pathways (**Fig. 6A, B**). KEGG pathway analysis showed that the DEGs were principally associated with the cytokine-cytokine receptor interaction and IL-17 signaling pathways (**Fig. 6C, 6D**).

**Figure 6.**
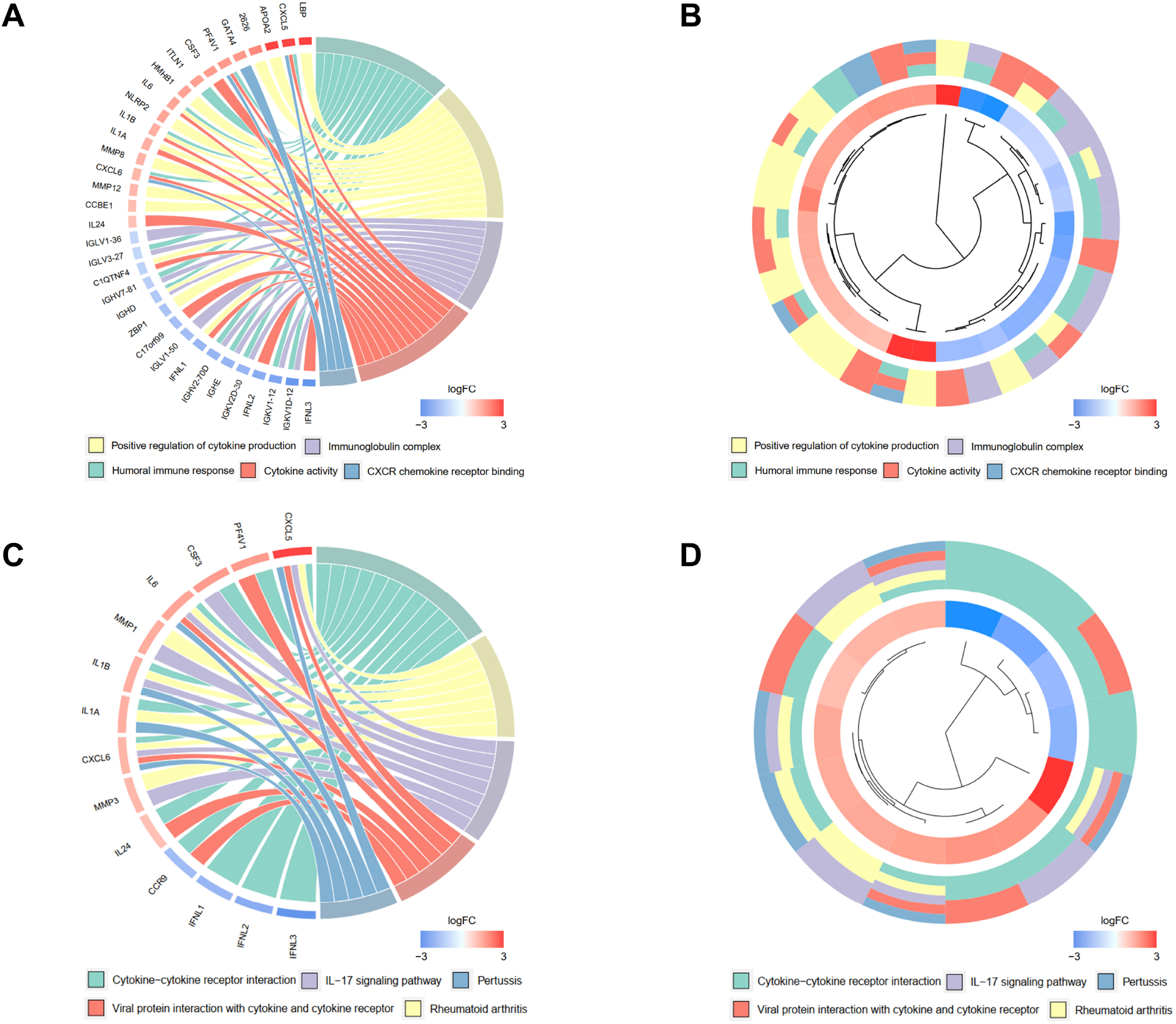
Functional enrichment analysis of differentially regulated genes (DEGs) between low- and high-risk groups. **A, B.** DEGs and pathways clustered by GO pathway enrichment analysis. **C, D.** DEGs and pathways clustered by KEGG pathway enrichment analysis.

### Exploration of association with immune status of EAC

To explore the correlation between the selected pyroptosis-related genes and gene-based signature with the immune microenvironment of EAC, analysis by TIMER database for each gene was initially performed. The results indicated that *ZBP1* expression was most significantly correlated with the infiltration signature of esophageal cancer, in which infiltrations of B cells (correlation coefficient = 0.366, *P* = 4.72e-07) and CD4+ T cells (correlation coefficient = 0.381, *P* = 1.41e-07) were with the most remarkable correlations (**Fig. 7A, Supplementary Fig. S1**). In addition, somatic copy number alterations of *ZBP1* were correlated with the infiltration levels of B cells, CD8+ T cells, macrophages and dendritic cells.

**Figure 7.**
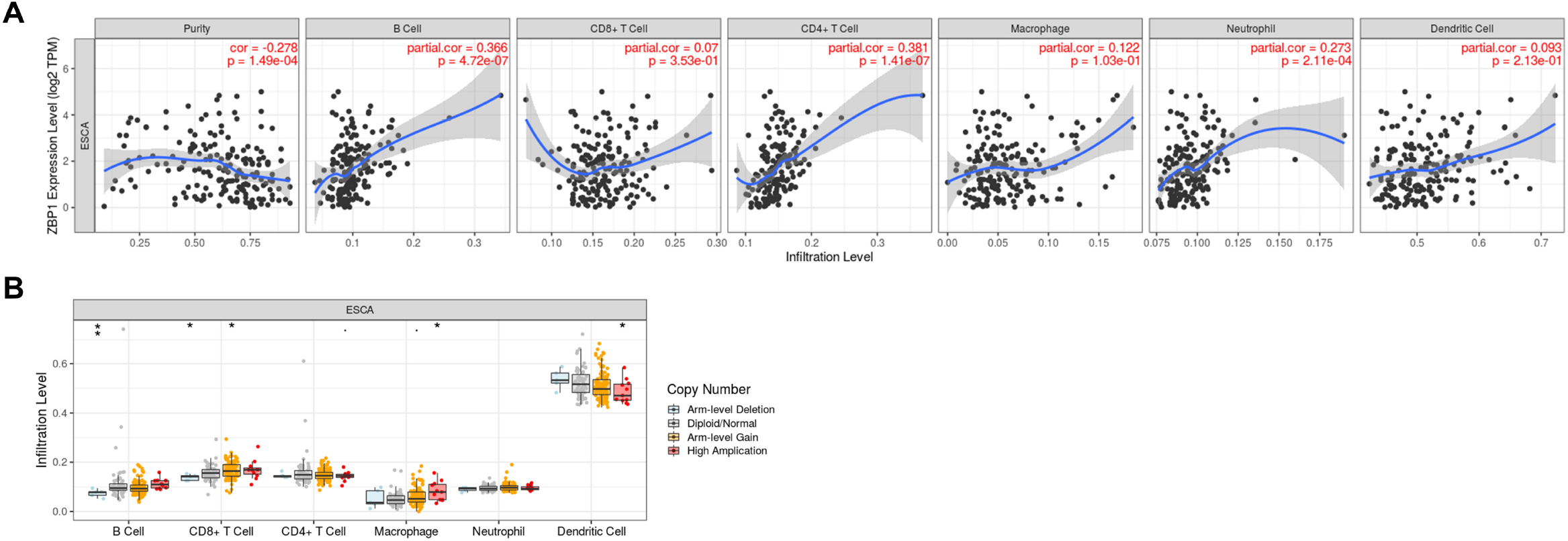
Correlations between immune cells and selected genes. **A.** ZBP1 and immune cells in esophageal cancer. **B.** Mutants (Arm-level Deletion, Arm-level Gain and High Amplication) of ZBP1 compared with Diploid/Normal in esophageal cancer.

The variations in the abundance of immune cell infiltration between low- and high-risk groups were further explored. The immune cells were analyzed in the TCGA (**Supplementary Table S3**) and GSE13898 cohorts (**Supplementary Table S4**). The overview of immune cell compositions was illustrated in **Fig. 8A** for the TCGA cohort, and **Fig. 8B** for the GSE13898 cohort. The high-risk group in the TCGA cohort possessed significantly higher infiltration levels of M2 macrophages, activated mast cells and eosinophils, whereas the infiltration levels of plasma cells were significantly lower (**Fig. 8C, 8D**). By contrast, the infiltration levels of memory B cells and M1 macrophages were upregulated in the high-risk group of the GSE13898 cohort, while those of naïve B cells were significantly downregulated (**Fig. 8E, 8F**).

**Figure 8.**
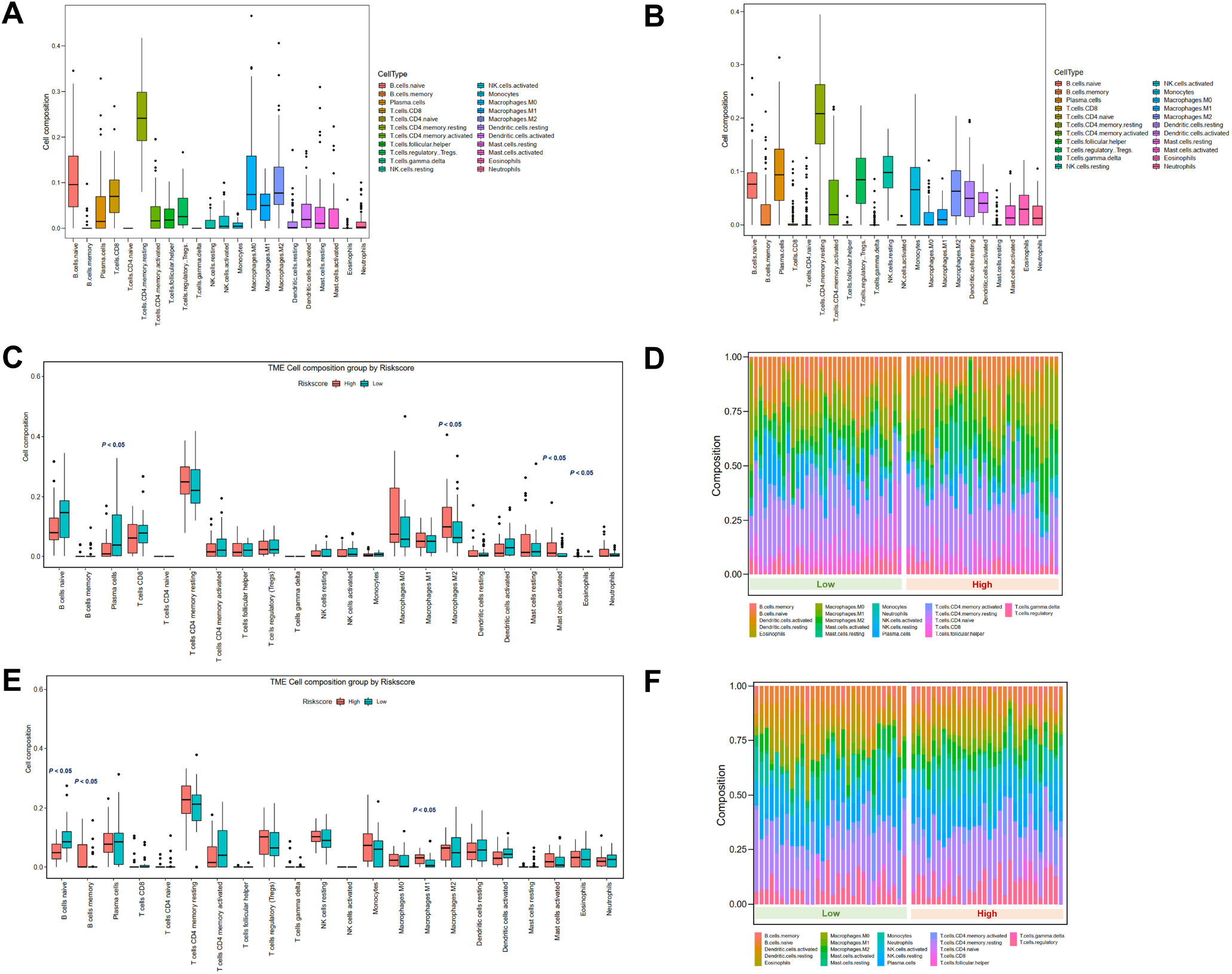
Tumor microenvironment immune cell composition in low- and high-risk groups. **A.** Overview of immune cell composition in TCGA cohort. **B.** Overview of immune cell composition in GSE13898 cohort. **C, D.** Differences in immune cell composition in TCGA cohort classified by risk groups by Wilcoxon test. **E, F.** Differences in immune cell composition in GSE13898 cohort classified by risk groups by Wilcoxon test.

## Discussion

Cell death serves as an essential barrier against the development of cancer, and pyroptosis is one of the major forms of programmed cell death ^32^. Mediated by the gasdermin family, pyroptosis is accompanied by immune and inflammatory responses ^7^. Emerging evidence has indicated the involvement of pyroptosis in cancer, while the dual role of pyroptosis has attracted the researchers’ attention ^7^. However, the role of pyroptosis in EAC remains largely unclear. In the present study, we comprehensively evaluated the pyroptosis-related gene profiles in EAC, and constructed a novel 5-gene risk signature (*CASP1, GSDMB, IL1B, PYCARD, ZBP1*) by LASSO Cox regression analysis. The 5-gene signature showed well performance for predicting EAC prognosis in both the internal and external validation cohorts. Further enrichment analyses revealed that the DEGs between the low- and high-risk groups were correlated with immune-related pathways. Tumor immune microenvironment analyses indicated that high-risk patients had decreased levels of infiltrating active immune cells and higher proportions of quiescent immune-cell infiltration.

Caspase 1 that encoded by *CASP1* is a member of the caspase family, which is activated by inflammasomes and induces pyroptosis ^33^. As tumor suppressor, the expressions of Caspase 1 are lower in various cancer tissues ^34^. Caspase 1 activation-mediated pyroptosis suppresses glioma cell growth and metastasis ^35^. Gasdermin B (GSDMB) belongs to the GSDM family and is more broadly expressed compared to other GSDM family members ^36^. The cleavage of GSDMB induced by lymphocyte-derived granzyme A triggers pyroptosis ^37^. By contrast, GSDMB is found highly expressed in several human cancer tissues, whereas the specific functions and mechanisms remain unclear ^38^. Interleukin 1 Beta (IL-1β) is a proinflammatory cytokine involved in pyroptosis. CASP-1 directly cleaves GSMD and precursor cytokines into pro-IL-1β, which initiates pyroptosis and maturation of IL-1β, respectively ^39^. IL-1β has pro-tumorigenic effects by promoting proliferation, migration and metastasis ^40^. Apoptosis-associated speck-like protein containing a CARD (ASC) is encoded by *PYCARD* gene and contains a caspase activation and recruitment domain (CARD) for binding and facilitating the activation of caspase 1 ^5^. The dual role of the inflammasome adaptor ASC is identified in cancer cells, as ASC has both pro-apoptotic and pro-inflammatory functions ^41^. Z-DNA-binding protein 1 (ZBP1)-NLR Family Pyrin Domain Containing 3 (NLRP3) is critical in inducing pyroptosis by leading to cytokine maturation and GSDMD cleavage ^42^. Deletion of *ZBP1* blocks tumor necroptosis and inhibits metastasis in breast cancer cells ^43^. However, the effects and mechanisms of *ZBP1* in tumors are poorly understood. In the current study, all of the 5 genes were upregulated in tumor samples compared to normal tissues, which is consistent with previous publications. Intriguingly, *GSDMB, PYCARD* and *ZBP1* were identified as down-regulated in the high-risk populations. It could be due to the pro-pyroptotic effect of GSDMB and the dual role of ASC in cancer cells. Therefore, pyroptotic-related genes might exert different effects in tumor development and progression depending on the biological context. Since the research of *ZBP1* in cancer is in its infancy, it is of importance for further exploration for its role and downstream-regulated pathways, especially for the involvement in the tumor microenvironment.

Crosstalks between the critical modes of programmed cell death exist, including apoptosis, pyroptosis and necroptosis, and part of their pathways are overlapping. For example, *CASP1, IL1B* and *PYCARD* are well-known regulators in the apoptotic pathways. It is inevitable that these pathways might have interactions as the tumor develops and progresses. Based on our enrichment analysis, immune response-correlated pathways are dysregulated in different risk groups, which indicates that inflammatory responses can be induced by pyroptotic cell death.

Nomograms are promising for use in clinical practice for evaluating the prognosis of EAC patients, in which the survival can be predicted using specific parameters. As indicated by the ROC curves, the nomogram demonstrates high predictive accuracy and sensitivity, especially for the prediction of 5-year survival. Compared to the conventional TNM staging and previously developed ferroptosis-related gene-signature in EAC ^44^, the pyroptosis-related gene signature-based nomogram, which integrates gene expression profiles and clinical parameters, more effectively predicts the prognosis of EAC patients. The use of nomogram based on integrated information can facilitate the prediction of prognosis, clinical decision-making and patient counseling ^45^.

Current studies on lymphocytes in tumor immunity predominantly focus on T cells, while the protective effect of B cells has also been revealed ^46^. By contrast, Mast cells have been reported to induce cancer growth ^47^. Activated T cells, natural killer cells and macrophages are potent suppressors that mediate tumor microenvironment and exert anti-tumor functions ^48, 49, 50^. Although some of the comparisons were not statistically different and might be contributed by the limited number of samples in both cohorts, accumulation of immune cells that promote cancer in the tumor microenvironment was generally observed in the high-risk group in both the TCGA and GEO cohort, while the compositions of tumor-protective immune cells were reduced compared to the low-risk group. In conclusion, the poor survival in the high-risk patients with EAC might be contributed by suppressed levels of anti-tumor immunity and alterations in the tumor microenvironment.

The strength of our study is that a systemic analysis was performed based on the TCGA and GEO cohorts, and the pyroptosis-related genes were assessed for the first time. Limitations also exist in our study. Due to the nature of publicly available dataset, the cases of EAC patients were limited, and the data of therapeutic responses were not available (e.g., chemotherapy, radiotherapy, immunotherapy). Specific mechanisms that pyroptosis-related genes regulate EAC development and progression remain to be explored. Large-scale and well-designed clinical studies for validation of our prediction model need to be performed. Despite the limitations, our study has provided a comprehensive overview of pyroptosis-related gene profiles in EAC.

In summary, we identified differentially expressed pyroptosis-related genes and developed a novel five-gene pyroptosis signature that significantly correlates with the survival of EAC patients. The pyroptosis-based signature is an independent prognostic factor and performs better than the TNM stage, which is promising for clinical application. Moreover, the genes in the pyroptosis-based signature might regulate anti-tumor immunity in tumor microenvironment, and further studies are warranted.

## Author Approvals

The authors have seen and approved the manuscript, and that it hasn’t been accepted or published elsewhere.

## Acknowledgement

The authors thank Prof. Ju-Seog Lee for kindly providing the data of the GSE13898 cohort.

## Data availability

The datasets are available in TCGA database (https://portal.gdc.cancer.gov) and GEO database (https://www.ncbi.nlm.nih.gov/gds/, accession number: GSE13898). R code used in this study are available from the corresponding author upon reasonable request.

## Supplementary Information

**Supplementary Figure S1. Correlations between immune cells and other genes in the pyroptosis signature.**

**Supplementary Table S1. Pyroptosis-related genes included in analysis.**

**Supplementary Table S2. Differentially expressed genes between the risk groups.**

**Supplementary Table S3. Immune cell composition in the TCGA cohort divided by risk groups.**

**Supplementary Table S4. Immune cell composition in the GSE13898 cohort divided by risk groups.**

